# Post-copulatory sexual selection is associated with sperm aggregate quality in *Peromyscus* mice

**DOI:** 10.1101/2020.02.08.939975

**Authors:** Kristin A. Hook, W. David Weber, Heidi S. Fisher

**Affiliations:** Department of Biology, University of Maryland, College Park, MD 20742, U.S.A.

**Keywords:** mating systems, sexual selection, sperm competition, sperm conjugation, sperm motility

## Abstract

In some species, sperm form coordinated groups that are hypothesized to improve their swimming performance in competitive contexts or to navigate through the viscous fluids of the female reproductive tract. Here we investigate sperm aggregation across closely-related species of *Peromyscus* mice that naturally vary by mating system to test the predictions that sperm aggregates (1) are faster than solitary sperm in species that females mate multiply to aid cells in sperm competition, and (2) outperform solitary sperm cells in viscous environments. We find significant variation in the size of sperm aggregates, which negatively associates with relative testis mass, a proxy for sperm competition risk, suggesting that post-copulatory sexual selection has a stabilizing effect on sperm group size. Moreover, our results show that sperm aggregates are faster than solitary sperm in some, but not all, species, and this can vary by fluid viscosity. Of the two species that produce the largest and most frequent groups, we find that sperm aggregates from the promiscuous *P. maniculatus* are faster than solitary sperm in every experimentally viscous environment but aggregation provides no such kinematic advantage under these same conditions for the monogamous *P. polionotus*. The reduced performance of *P. polionotus* aggregates is associated with less efficient aggregate geometry and the inclusion of immotile or morphological abnormal sperm. Our cross-species comparison yields insight into the evolution of sperm social behaviors, provides evidence of extensive variation in the *Peromyscus* lineage, and reveals that differences in sperm aggregate quality associate with post-copulatory sexual selection.

## Introduction

Sperm cells are one of the most diverse cell types in nature and exhibit striking variation both within and across taxa (Pitnick et al. 2009). In addition to being morphologically diverse, sperm may exhibit complex, emergent behaviors, including sperm conjugation, which occurs when two or more cells join together for motility or transport through the female reproductive tract before dissociating prior to fertilization (Pitnick et al. 2009; Higginson and Pitnick 2011; Schoeller et al. 2020). Although this phenomenon is relatively rare, sperm aggregation has evolved multiple times across independent lineages of internally fertilizing species (reviewed in Immler 2008; Pitnick et al. 2009; Higginson and Pitnick 2011; Umezu et al. 2020), suggesting that these unique sperm-sperm interactions may be a convergent adaptation driven by a shared selective force.

The primary selective force that has been hypothesized to drive the evolution of sperm aggregation is post-copulatory sexual selection and, more specifically, sperm competition (Moore et al. 2002; Immler et al. 2007; Fisher and Hoekstra 2010). Sperm competition occurs when females mate with multiple males within a reproductive cycle (i.e., polyandry), allowing the sperm produced by those males to compete for fertilization of her ova (Parker 1970; Simmons 2001). Within these competitive contexts, collective sperm groups are posited to be advantageous if the combined force generated by their multiple flagella enable them to swim faster (Moore et al. 2002; Immler et al. 2007) or to migrate more efficiently through the viscous or viscoelastic secretions of the female reproductive tract (Moore and Taggart 1995; Suarez 2016), thus enhancing their fertilization success (Birkhead et al. 1999; Gage et al. 2004). Among the relatively few studies that have attempted to elucidate the advantages of collective sperm behavior, only some have provided empirical support that sperm aggregates are faster than solitary cells in competitive or complex environments (Hayashi 1998; Moore et al. 2002; Fisher and Hoekstra 2010; Pearcy et al. 2014; but see Ishijima et al. 1999). However, given the mixed results yielded by interspecific analyses (i.e., Moore and Taggart 1995; Immler et al. 2007; Pitnick et al. 2009; Woolley et al. 2009; Tung et al. 2017), the benefits of sperm conjugation are likely to be taxon- and context-dependent.

One potential reason for inconsistencies regarding the kinematic benefits of sperm aggregates across taxonomic groups is that they vary in when and how they form (Higginson and Pitnick 2011; Monclus and Fornes 2016) and, consequently, often differ in the number of cells that are present within them (Pizzari and Foster 2008). For example, some mammalian sperm conjugates are derived from spermatogenic processes and assemble during epididymal transport (Higginson and Pitnick 2011), forming groups with very little intraspecific variation; these sperm are often molecularly “glued” to one another, such as the bi-flagellate pairs in grey short-tailed opossums (*Monodelphis domestica*, Taggart et al. 1993), sperm ‘rouleaux’ of five or more cells in guinea pigs (*Cavia porcellus*, McGlinn et al. 1979), or bundles of roughly 100 cells in monotremes (Nixon et al. 2016). Conversely, some mammalian sperm conjugates are derived from post-spermatogenic processes and assemble after ejaculation (Higginson and Pitnick 2011), thus forming more variably-sized groups. For example, sperm may form temporary clusters of up to sixteen cells in bulls (*Bos Taurus*, Tung et al. 2017) or more fixed groups of 3-30 cells in house mice (Immler et al. 2007), 5-50 cells in the Norway rat (Immler et al. 2007), or hundreds to thousands of cells in the wood mouse (*Apodemus sylvaticus*, Moore et al. 2002). These systems with post-spermatogenic aggregation are ideal for assessing the kinematic consequences of aggregation because their ejaculates contain both solitary and aggregated sperm cells, thereby enabling a direct comparison of these naturally occurring sperm types produced by the same male. Furthermore, these systems allow for investigations into the functional significance of aggregate size variation upon which selective pressures like sperm competition can act.

Closely-related species of mice in the genus *Peromyscus* naturally vary in their mating systems (Turner et al. 2010; Bedford and Hoekstra 2015), and at least two species are known to produce sperm cells that form temporary, motile groups after ejaculation (Fisher and Hoekstra 2010). Of these, sperm produced by the promiscuous deer mouse (*P. maniculatus*), selectively group with the most closely-related sperm to form aggregates of up to 30 cells (Fisher and Hoekstra 2010), but there is a non-monotonic association between aggregate size and swimming velocity, indicating that only aggregates of a certain size are faster than solitary cells (Fisher et al. 2014). In comparison, sperm from its monogamous congener, the oldfield mouse (*P. polionotus*), indiscriminately group with both related and unrelated cells (Fisher and Hoekstra 2010), and their aggregates are less likely to be optimally-sized (Fisher et al. 2014). These studies provide compelling evidence that post-copulatory sexual selection has influenced the evolution of sperm aggregation within this system, but the broader extent of sperm aggregation within the *Peromyscu*s lineage remains unknown.

In this study, we examine the function of sperm aggregation and the role of post-copulatory sexual selection in shaping the evolution of these collective sperm behaviors in *Peromyscus* mice. We compared sperm from three monogamous species (*P. californicus, P. eremicus, P. polionotus*) and three species in which females mate with multiple males (*P. maniculatus, P. leucopus, P. gossypinus*), under consistent, controlled *in vitro* conditions that we experimentally altered to modify the viscosity of the fluid. From these observations, we (1) describe the naturally occurring intra- and inter-specific variation in sperm aggregation in these species, (2) test the prediction that sperm aggregates are faster than solitary cells in the species with more intense sperm competition, and (3) test the prediction that sperm aggregates are faster than solitary cells in viscous environments.

## Materials and Methods

### (a) Sperm collection and standardization

We obtained captive *Peromyscus maniculatus bairdii, P. polionotus subgriseus, P. leucopus, P. eremicus*, and *P. californicus insignis* from the Peromyscus Genetic Stock Center at the University of South Carolina, and *P. gossypinus gossypinus* from Dr. Hopi Hoekstra at Harvard University and housed them in same-sex cages at 22°C on a 16L:8D cycle in accordance with guidelines established by the Institutional Animal Care and Use Committee at the University of Maryland in College Park (protocol R-Jul-18-38). We sought samples from all available captive *Peromyscus* species, avoiding wild-caught specimens to control for variation due to life experience. All males were sexually mature and we accounted for relatedness by assigning siblings a unique ‘Family’ ID.

To collect sperm, we euthanized males via isoflurane overdose and cervical dislocation, recorded their body and testes weight, and then removed a single caudal epididymis. We made several small incisions in the epididymis and submersed it in a centrifuge tube containing medium (Modified Sperm Washing Medium, Irvine Scientific, USA) that was pre-warmed at 37°C. To help standardize sperm concentrations across males based on qualitative assessments of the epididymal tissue size, we varied the media volume (50µl − 1000µl) by male and accounted for these volumetric differences when estimating sperm counts. We agitated the tube containing tissue and media at 300rpm (ThermoMixer F1.5, Eppendorf, Germany) at 37°C for ten minutes, inverting it at the five- and ten-minute marks and then incubating it for two minutes undisturbed. Using pipette tips with a cut tip to create a wider opening and avoid breaking up aggregates, we collected live sperm for analysis from just below the meniscus of the solution to enrich for the most motile sperm (Magdanz et al. 2019). We used a computer-assisted sperm analysis (CASA) system (Ceros II^™^ Animal, Hamilton Thorne, USA) to rapidly assess sperm concentrations within each sample, which we later verified via manual sperm counts using a hemocytometer. We then diluted samples as needed with pre-warmed sperm medium to reach a standard concentration of cells across all samples (Supplementary Materials).

### (b) Live sperm analysis

To prepare live sperm samples for *in vitro* observations in a ‘low-viscosity’ environment, we gently reverse pipetted 4µl of the sperm solution into 12µl of pre-warmed medium on a plastic slide within a 9mm x 0.12mm imaging spacer (Grace Bio-Labs, USA) and covered it with a plastic cover slip (Supplementary Materials). To prepare the same sperm samples for *in vitro* observations in a ‘high-viscosity’ environment, we mixed 4µl of sperm solution with 12µl of pre-warmed medium enriched with methylcellulose (Sigma Aldrich M 7140; 15cP, 2% in water; Suarez and Dai 1992) to produce three experimental treatment groups with increasing viscosity levels of 0.75%, 1.5%, or 2.25% methylcellulose solutions (see Supplementary Materials). We recorded a minimum of five 5-second videos at 60 frames/second per male, except in one case in which we were only able to obtain four videos. The mean (±SE) duration of time between tissue dissection and video observations (i.e., ‘post-harvest time’) was 59 (±1) minutes, which varied slightly depending on how long it took to properly dilute each sample.

From the sperm videos obtained using the CASA system, we manually scored the number of sperm cells present within each track (i.e., aggregate size) by observing at least three different frames per track. When tracking sperm, CASA automatically excluded any tracks in which cells left the field within the first 10 frames or entered the field or started moving after the first 10 frames. Through our manual evaluation of each track, we further removed tracks if they involved cells or aggregates that were attached to debris or the slide surface, too out of focus to count the number of cells, interacted with other cells or aggregates (e.g., colliding, joining, separating, or impeding movement), or had a distance average path < 30 µm (often repeats of longer, more informative tracks). From the remaining tracks, we categorized each track as either “solitary” or “aggregated”. We calculated mean *sperm aggregate size* (i.e., sperm cells per aggregate) by dividing the number of aggregated sperm cells by the total number of aggregates. We further scored aggregates that were comprised of at least one cell that was immotile (i.e., unmoving or stuck), misaligned (i.e., cells opposed one another, interacted loosely, or formed via attachment with the flagella rather than at the head region), or morphologically abnormal (i.e., kinked at the midpiece or missing a head or flagella) to determine the proportion of aggregates that featured these abnormalities (‘abnormal’) compared to those that did not (‘normal’). We pooled these data either across all tracks per male or across all males per species to characterize sperm aggregates by males and species, respectively (Crawley 2013). Last, we calculated the coefficient of variation (CV) for aggregate size within each species using the following formula: (standard deviation/mean) x 100.

From the same CASA video recordings used to calculate the size and composition of sperm aggregates, we collected swimming performance data for all motile sperm tracks. To create a dataset of only motile sperm tracks, we added additional restrictions to those outlined above for our aggregate analysis, including the removal of cells and aggregates that were immotile (i.e., unmoving) or poorly tracked by CASA (i.e., featuring a low number of detection points, points that jumped suddenly, or points that picked up other regions of the cell). We additionally removed any solitary cells that were morphologically abnormal (e.g., featuring a kinked midpiece section that resulted in a backwards swimming pattern). From these data, we determined the speed (curvilinear velocity, VCL) for both solitary and aggregated sperm cells for each male by pooling their motile tracks within the different media treatment groups (i.e., low-viscosity or high-viscosity).

### (c) Statistical Analyses

We performed all statistical analyses using R version 3.4.2 (R Core Team 2016) and created all figures using the ‘ggplot2’ package with R (Wickham 2016). To validate model normality, we visually inspected diagnostic plots (qqplots and plots of the distribution of the residuals against fitted values). Only the best fitting models are reported here.

To investigate differences in sperm aggregate size across our focal *Peromyscus* species, we used a linear model (LM) and mean values for each male. We excluded one *P. californicus* male whose measurement represented a clear outlier; the removal of this individual did not qualitatively change the results (see Supplementary Materials). We initially used a linear mixed model (LMM) using the lmer function from the “lme4” R package (Bates et al. 2015) with ‘Family ID’ as a random factor, but this variable did not significantly contribute to the residual variability in the response variable. Predictors that were considered for the initial LMM included male age, pairing status, post-harvest time, total sperm cells, number of recorded videos, and the ratio of sperm cells per video. We considered predictors with *p*-values < 0.20 for the final model, but first screened each for collinearity with other significant predictors using a linear model. Whenever collinearity was present, only the predictor with the greater relative significance was included in the model. We dropped non-significant explanatory variables one at a time based on model comparisons using an analyses of variance test to determine the minimal adequate model, which included total sperm cells and species as predictors. We performed post-hoc pairwise species comparisons using Tukey HSD adjustments for multiple comparisons from the “LSmeans” R package (Lenth 2016).

We next examined the relationship between mean sperm aggregate size and sperm aggregate size CV with the mean relative testis size using mean values for each species. We accounted for the evolutionary relationships among the species used in this study by adopting a phylogenetic generalized least squares (PGLS) approach (Pagel 1999; Freckleton et al. 2002) using the “caper” (Orme et al. 2013) and “APE” (Paradis et al. 2004) packages in R. To do so, we used an ultra-metric phylogenetic tree of *Peromyscus* (provided by Dr. Roy Neal Platt II, Texas Biomedical Research Institute) based on sequence variation in the mitochondrial gene, cytochrome B, as a covariate in regression analyses. This phylogeny was similar to other previously established *Peromyscus* phylogenies (Bradley et al. 2007; Turner et al. 2010). Predictors that were considered for both regression models included testis mass and body mass – a method better suited to estimating relative testis weight rather than using the ratio of testis to body mass or residuals (GarcÍa-Berthou 2001; Lüpold et al. 2009).

To test whether sperm aggregates offer a kinematic benefit, we used a paired student’s t-test for each species separately and mean values per male to control for inter-male variation; we compared the VCL of sperm aggregates to the VCL of solitary sperm cells. We performed these statistical tests for motility data collected when sperm were suspended in either low- or high-viscosity media to examine whether the kinematic benefits of aggregation are context dependent.

To compare the composition of sperm aggregates, we used the number of abnormal aggregates and total sperm aggregates summed across all males within each species (Crawley 2013) within a multiple proportions test to examine whether species differ in the proportion of sperm aggregates that feature abnormalities. We then compared the VCL of normal sperm aggregates to abnormal sperm aggregates using paired student’s t-test for each species separately and mean values per male to control for inter-male variation.

## Results

We found that sperm aggregate size differs among *Peromyscus* species (LM: F_6,126_ = 56.31, *p* < 0.001; Figure 1A, Table 1), indicating substantial trait variation within this lineage. We found greater variance in the number of aggregated cells between than within species (proportion: s^2^ across species = 1.96; s^2^ within species < 1.00, except for *P. polionotus* [s^*2*^ = 2.15]; Figure S1). Pairwise comparisons adjusted for multiple comparisons using LSmeans revealed that *P. gossypinus, P. leucopus*, and *P. californicus* produce the smallest aggregates (*p* > 0.05 for pairwise comparisons), the latter of which produces sperm cell aggregates that are statistically similar in size to those produced by *P. eremicus* (*p* = 0.92; Figure 1A). Conversely, the largest aggregates were observed in two sister species, *P. maniculatus* and *P. polionotus*, which produced similarly sized aggregates that did not statistically differ (*p* = 0.88).

**TABLE 1.** Fixed effects from a linear model examining differences in sperm aggregate size across focal *Peromyscus* species.

| Model Term | Beta (SE) | Exp (beta) | 95% CI | *t* | Pr(>\|z\|) |
| --- | --- | --- | --- | --- | --- |
| Intercept | 3.90 (0.26) |  |  |  |  |
| Total Sperm Cells | 0.00 (0.00) | 0.50 | (0.50, 0.50) | 5.82 | < **0.001** |
| *P. eremicus* | -1.97 (0.24) | 0.12 | (0.08, 0.18) | -8.14 | < **0.001** |
| *P. gossypinus* | -2.77 (0.24) | 0.06 | (0.04, 0.09) | -11.53 | < **0.001** |
| *P. californicus* | -2.19 (0.23) | 0.10 | (0.07, 0.15) | -9.67 | < **0.001** |
| *P. leucopus* | -2.68 (0.24) | 0.06 | (0.04, 0.10) | -11.18 | < **0.001** |
| *P. polionotus* | 0.04 (0.23) | 0.51 | (0.40, 0.62) | 0.15 | 0.88 |

All rows are compared with the intercept – *P. maniculatus*. 95% confidence intervals (CI) were calculated for each effect size.

**Figure 1.**
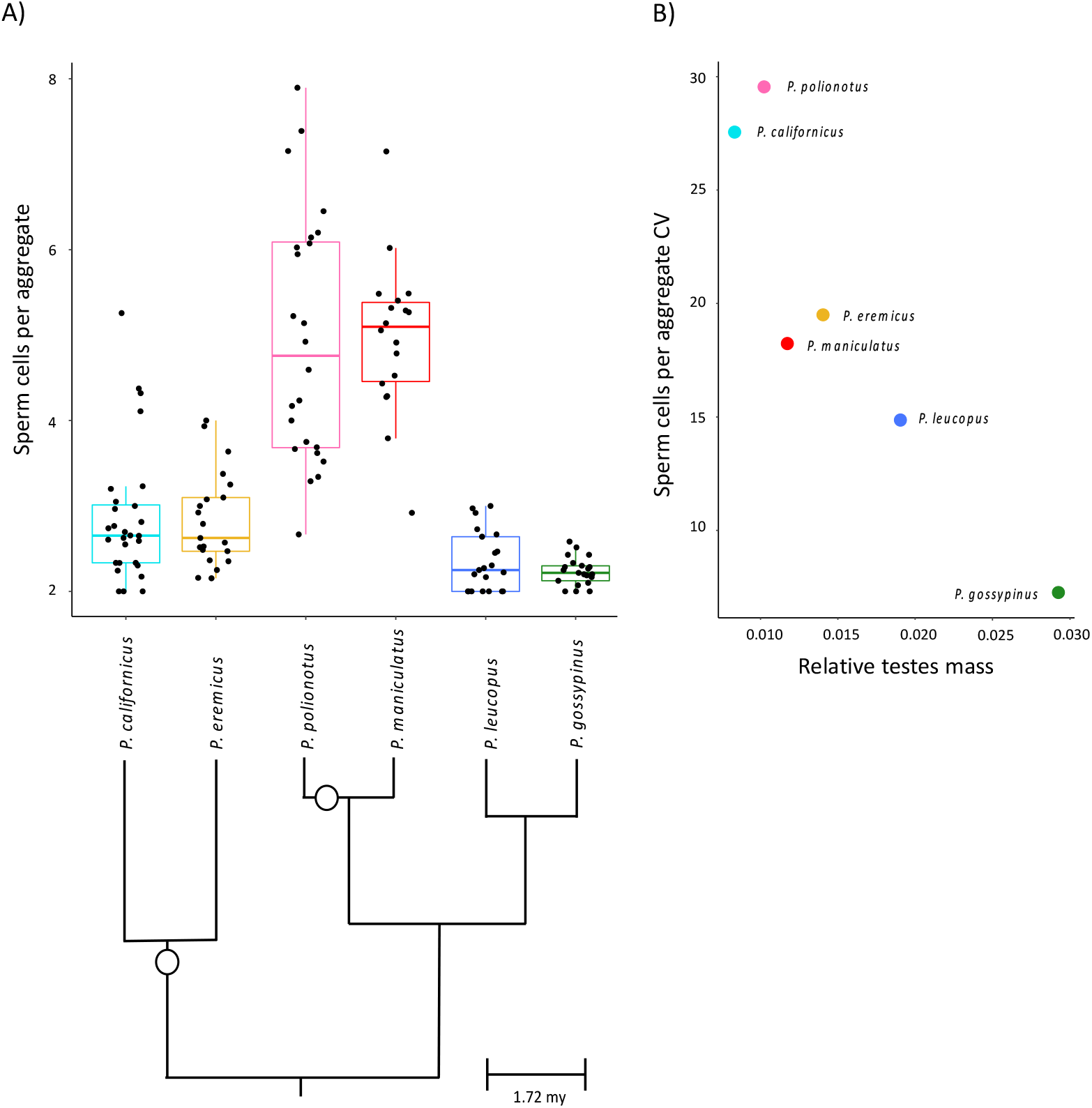
Sperm aggregate size differs within and among *Peromyscus* mice, and variability in this trait negatively associates with relative testis size. (A) Observed variation in the number of sperm cells per aggregate produced by males under controlled, *in vitro* conditions. Box-plots represent median and interquartile ranges with mean values per male overlaid as black dots. Species relationships are indicated within the phylogeny (adapted from Bradley et al. 2007), and open circles represent the evolutionary transitions to monogamy (Bedford & Hoekstra 2015). (B) When controlling for phylogenetic relationships across these species using a PGLS model, the coefficient of variation (CV) for the number of sperm cells in aggregate negatively correlates with relative testes mass. Note truncated y-axes.

Moreover, we found a negative association between the coefficient of variation (CV) for sperm aggregate size and testis weight (PGLS: F_2,3_= 6.96, *p* = 0.034) across these focal species (Figure 1B, Table 2), although there was no such association between aggregate size itself and testis weight (PGLS: F_2,3_= 3.058, *p* = 0.153; Table 2). The sperm aggregate size CV for each species were as follows: 27.6% *P. californicus*, 19.5% *P. eremicus*, 29.5% *P. polionotus*, 18.2% *P. maniculatus*, 14.9% *P. leucopus*, and 7.3% *P. gossypinus*.

**TABLE 2.** Results from PGLS models examining the number of cells in aggregate or the coefficient of variation (CV) of this trait, in relation to relative testis mass for all focal *Peromyscus* species

| Response | Predictor | *t* | *P* | *R*<sup>2</sup> | $\lambda$ |
| --- | --- | --- | --- | --- | --- |
| Number of cells in aggregate | Testis mass (mg) | -1.9032 | 0.153 | 0.45 | 0.000<br>(1.00, 0.50) |
|  | Body mass (mg) | -1.0037 | 0.389 |  |  |
| Number of cells in aggregate CV | Testis mass (mg) | -3.7267 | **0.034** | 0.70 | 0.000<br>(1, 0.01) |
|  | Body mass (mg) | 1.1829 | 0.322 |  |  |

In all models, the branch length transformations for lambda, $\lambda$ , were set using maximum likelihood ('ML'). Lower and upper bounds for this term are noted within parentheses.

When comparing the mean curvilinear velocity (VCL) of aggregated to solitary sperm cells, we found that sperm aggregates are significantly faster in all experimental media (0%, 0.75%, 1.5%, or 2.25% methylcellulose) in only one species, the promiscuous deer mouse (*P. maniculatus*; Table 3, Figure 2, Table S1). In the low-viscosity environment (0% methylcellulose), we found that sperm aggregates are slower in two species, *P. polionotus* and *P. gossypinus*; the speed of sperm aggregates did not statistically differ from solitary cells in *P. californicus, P. eremicus*, and *P. leucopus* (Table 3, Figure 2). Within all of the high-viscosity environments (0.75%, 1.5%, or 2.25% methylcellulose), we found that sperm aggregates were faster that solitary sperm in *P. californicus* (Table 3, Figure 2, Table S1). We also found that for at least one higher viscosity environment, sperm aggregates were relatively faster than solitary sperm in *P. eremicus* (1.5% methylcellulose), *P. gossypinus* (1.5% methylcellulose), and *P. leucopus* (2.25% methylcellulose); otherwise, there were no significant differences between the speed of sperm aggregates and solitary sperm cells (Table 3, Figure 2, Table S1).

**TABLE 3.** Results from an intra-male analysis for all focal *Peromyscus* species comparing the curvilinear velocity (VCL) of solitary and aggregated sperm in low- and high-viscosity environments to test whether sperm aggregates confer a kinematic advantage (shaded in gray)

| *Peromyscus*<br>Species | Low Viscosity (0% methylcellulose) |  |  |  |  | High Viscosity (0.75% methylcellulose) |  |  |  |  |
| --- | --- | --- | --- | --- | --- | --- | --- | --- | --- | --- |
| | Mean (±SE)<br>VCL ($\mu\text{m}/\text{sec}$) | | Student's paired t-test | | | Mean (±SE)<br>VCL ($\mu\text{m}/\text{sec}$) | | Student's paired t-test | | |
|  | Solitary<br>cells | Aggregated<br>cells | *t* | df | *p* | Solitary<br>cells | Aggregated<br>cells | *t* | df | *p* |
| *californicus* | 142.3 ± 3.8 | 144.4 ± 4.2 | -1.0545 | 28 | 0.301 | 81.0 ± 8.0 | 93.0 ± 8.0 | -3.9495 | 9 | < **0.01** |
| *eremicus* | 123.4 ± 2.6 | 114.3 ± 4.7 | 1.7308 | 20 | 0.099 | 91.3 ± 3.9 | 94.6 ± 5.9 | -0.7174 | 9 | 0.491 |
| *polionotus* | 160.4 ± 4.5 | 133.1 ± 3.3 | 9.4575 | 23 | < **0.001** | 100.6 ± 6.6 | 93.9 ± 5.2 | 2.0349 | 13 | 0.063 |
| *maniculatus* | 163.5 ± 4.7 | 177.7 ± 6.8 | -2.2482 | 17 | **0.038** | 115.4 ± 4.3 | 126.5 ± 5.6 | -2.9397 | 11 | **0.014** |
| *leucopus* | 129.8 ± 2.5 | 128.6 ± 2.9 | 0.24811 | 20 | 0.807 | 101.6 ± 3.4 | 100.8 ± 5.0 | 0.3243 | 9 | 0.753 |
| *gossypinus* | 106.5 ± 2.7 | 90.1 ± 3.2 | 5.4048 | 20 | < **0.001** | 89.1 ± 4.6 | 90.4 ± 5.8 | -0.6384 | 10 | 0.538 |

**Figure 2.**
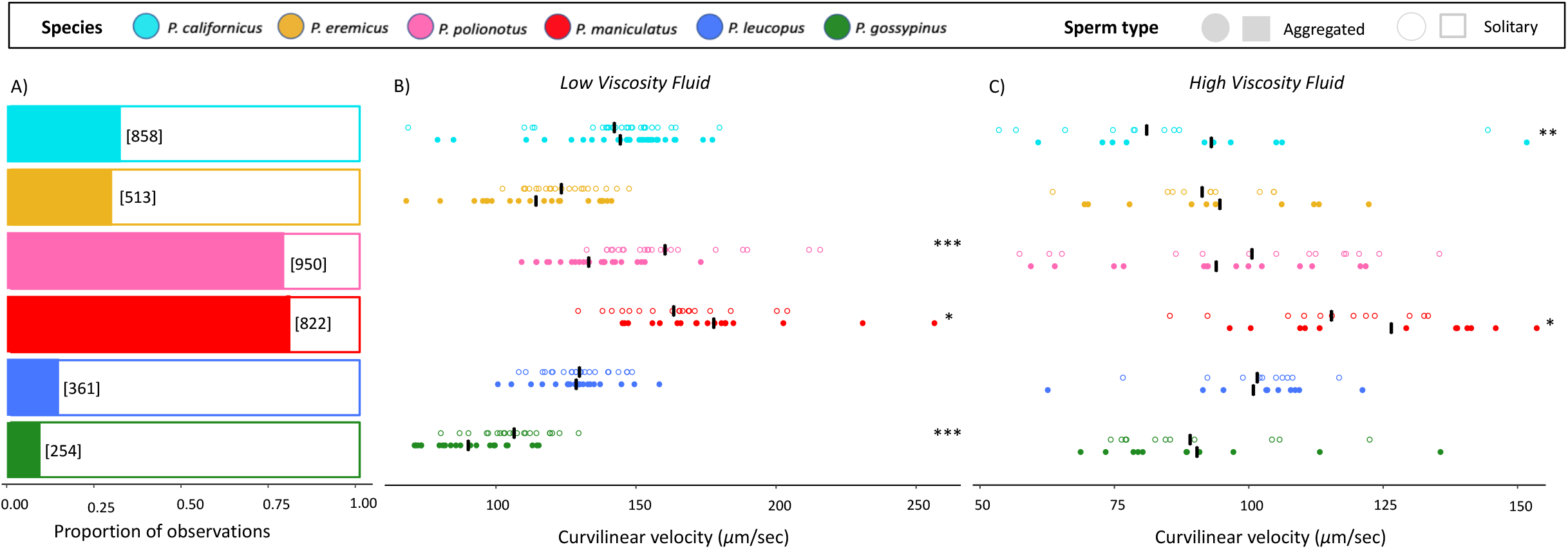
The frequency of sperm aggregates varies across focal *Peromyscus* species and can affect sperm swimming performance. (A) Sperm aggregates (solid bars) and solitary sperm (open bars) are simultaneously produced by males of all focal species, allowing for a direct comparison of their speeds while controlling for inter-male variation to test the hypothesis that sperm aggregates provide a competitive advantage. The summed number of sperm aggregates observed within each species is indicated in brackets. The curvilinear velocity of aggregated (closed circles) and solitary sperm (open circles) observed in either low (B) or high (C) viscosity media to test the hypothesis that sperm aggregates are faster in viscous environments. In both environments, sperm aggregates are faster in *P. maniculatus*. Aggregates are also faster in *P. californicus* in the high-viscosity environment, but aggregates are slower in *P. polionotus* and *P. gossypinus* in the low-viscosity environment. All other species produced aggregates that did not differ in speed from solitary sperm. Circles represent mean values per male, and black dashes represent species mean per sperm type. * *p* < 0.05, ** *p* < 0.01, *** *p* < 0.001. Note the truncated x-axes.

When comparing the composition of sperm aggregates, we observed the highest number of abnormal aggregates in the three species with promiscuous mating systems (*P. maniculatus* =107, *P. leucopus* = 128; *P. gossypinus* = 58) and the lowest number of abnormal aggregates in the three species with monogamous mating systems (*P. eremicus* = 266, *P. polionotus* = 273, *P. californicus* = 187; Figure 3a). When comparing abnormal aggregates to the total number of aggregates observed, we found that species differ in the proportion of ‘abnormal’ aggregates that they produce (multiple proportions test: *χ*^2^ = 270.12, df = 5, *p* < 0.001), with *P. maniculatus* producing the lowest proportion of abnormal sperm aggregates (abnormal/total: 107/822) and *P. eremicus* producing the highest proportion of abnormal aggregates (266/513) compared to all other species (*P. californicus* 187/858; *P. polionotus* 273/950; *P. leucopus* 128/361; *P. gossypinus* 58/254; Figure 3a). Importantly, we found that abnormality can negatively impact aggregate speed; abnormal sperm aggregates have a significantly lower VCL compared to normal sperm aggregates in all species except *P. leucopus* and *P. gossypinus* (Table 4, Figure 3b).

**TABLE 4.** Results from an intra-male analysis for all focal *Peromyscus* species comparing the curvilinear velocity of sperm aggregates featuring one or more cells that were immotile, misaligned, or morphologically flawed with aggregates that lacked abnormalities

| *Peromyscus*<br>Species | Student's paired t-test |  |  |
| --- | --- | --- | --- |
|  | *t* | df | *p* |
| *californicus* | 3.967 | 27 | < **0.001** |
| *eremicus* | 2.816 | 19 | **0.011** |
| *polionotus* | 4.996 | 20 | < **0.001** |
| *maniculatus* | 5.354 | 15 | < **0.001** |
| *leucopus* | 0.807 | 14 | 0.433 |
| *gossypinus* | 1.165 | 17 | 0.260 |

**Figure 3.**
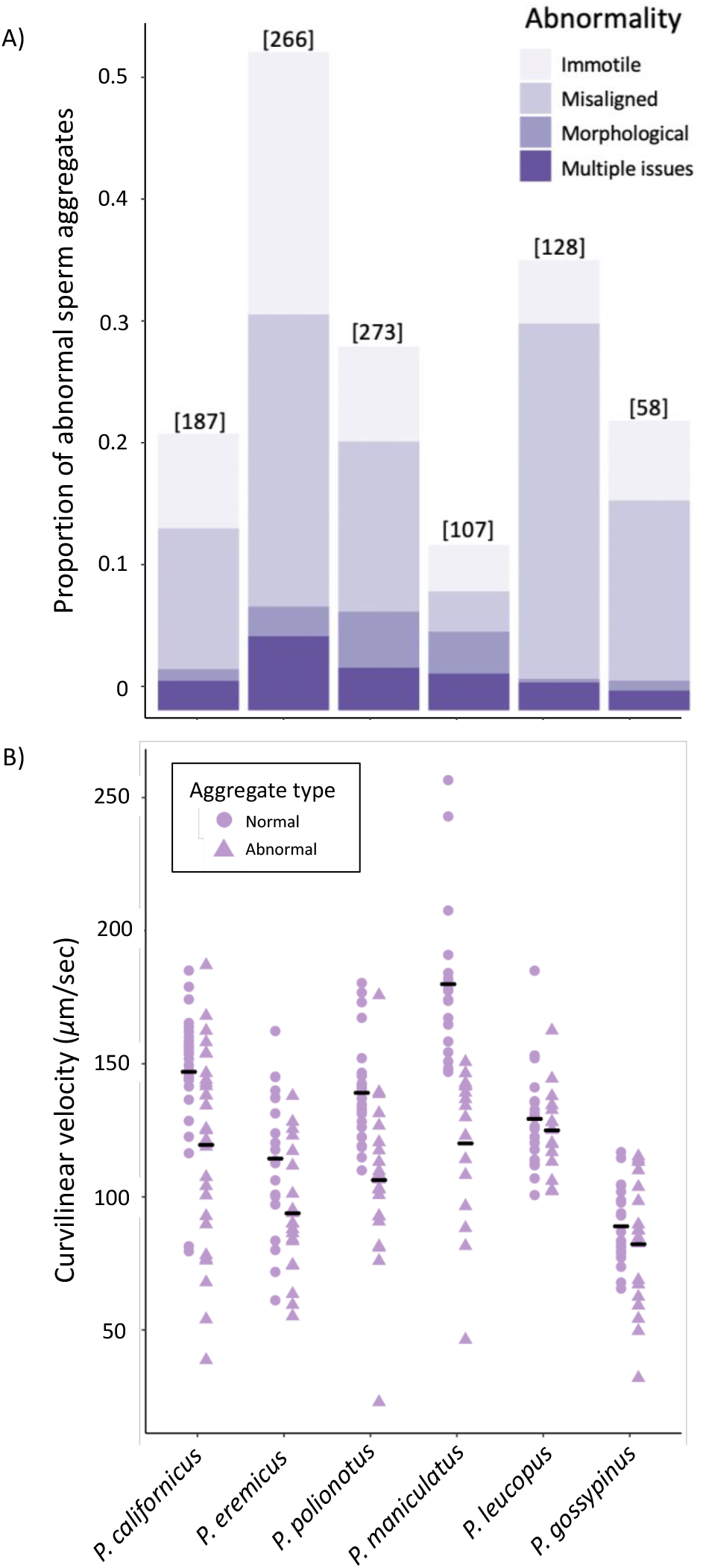
The alignment of sperm within aggregates and the inclusion of abnormal or immotile cells varies across focal *Peromyscus* species and impacts aggregate speed. (A) Sperm aggregates were scored as ‘abnormal’ if they featured one or more cells that were immotile, misaligned, morphologically abnormal, or had a combination of these issues. The total number of abnormal aggregates observed per sperm are indicated in brackets. (B) Abnormal aggregates (triangles) are slower than normal aggregates (circles) in all but two species (*P. leucopus* and *P. gossypinus*). Symbols represent mean values per male, and black dashes represent species mean per aggregate type.

## Discussion

Collective sperm behaviors have evolved independently in a number of taxa (Higginson and Pitnick 2011; Schoeller et al. 2020), yet their functional significance remains poorly understood.

Here we examine the evolution of sperm aggregation in closely-related species of *Peromyscus* mice that vary by mating strategy. In *Peromyscus*, males simultaneously produce solitary and aggregated sperm cells, allowing us to directly compare these naturally occurring sperm types. These comparisons revealed interspecific differences in the size of sperm aggregates that form, and importantly, that variability in aggregate size negatively associates with relative testis weight, a robust proxy for level of sperm competition in rodents (Ramm et al. 2005), suggesting that this trait may be under stabilizing selection. We also found that collective sperm groups enhance motility in some, but not all, species and in some, but not all, environments. Our data revealed that these differences in aggregate speed are associated with the orientation and composition of the cells within the group (Fisher et al. 2014), and that those species that evolved under high levels of sperm competition produced the fewest number of abnormal aggregates. Together, our findings suggest that post-copulatory sexual selection can play an important role in shaping overall sperm aggregate quality, including motility, but that relaxed selection in monogamous taxa may have enabled less efficient strategies to persist (van der Horst and Maree 2014), thereby generating diversity in collective sperm behaviors within and between closely-related species.

Under controlled *in vitro* conditions, we found distinct differences in the size of sperm aggregates produced by males across six closely related species of *Peromyscus* mice, with two sister species (*P. maniculatus* and *P. polionotus*) forming large sperm aggregates, two sister species (*P. californicus* and *P. eremicus*) forming moderately sized aggregates, and two sister species (*P. leucopus* and *P. gossypinus*) forming small sperm aggregates. Moreover, we found that that species with relatively larger testes, which positively associates with sperm competition risk (Ramm et al. 2005), exhibit less variation in aggregate size. This result supports the prediction that relaxed sperm competition allows for greater intermale variation to persist in a population (Calhim et al. 2007) and suggests that post-copulatory sexual selection may be stabilizing sperm aggregate size for a species-specific ‘optimum’ (Fisher et al. 2014). Other studies have shown that the strength of sexual selection regulates variance in sperm morphology across taxa and at multiple levels of organization, including within- and between-males (Immler et al. 2008; Fitzpatrick and Baer 2011; Carballo et al. 2019) and within- and between-species (Calhim et al. 2007; Rowley et al. 2019). Similarly, a study on sperm bundles across ten *Carabus* ground beetles also found intense selection on bundle size, which are dimorphic and either small or large; the large, but not small, sperm bundles are positively correlated with measures of sperm competition risk, including copulatory piece length and mate guarding, suggesting that diversity of large sperm bundles is associated with sexual selection (Takami and Sota 2007). In contrast to these findings that sperm competition drives sperm-sperm interactions, a study on the evolution of these interactions in diving beetles found that variation in sperm conjugation associates more with female reproductive tract architecture (Higginson et al. 2012). Therefore, while our results show that the uniformity of sperm aggregate size associates with an increase in sperm competition risk, mechanisms of female control (Eberhard 1996) may also play an important evolutionary role as a stabilizing force on this trait.

We found that aggregation enhances sperm motility in some, but not all, species and in some, but not all, environments; thus, our results provide mixed support for the hypotheses that the combined force generated by an aggregate’s multiple flagella improves motility in the context of sperm competition (Moore et al. 2002; Immler et al. 2007) and/or may allow them to migrate more efficiently through the viscous or viscoelastic secretions of the female reproductive tract (Moore and Taggart 1995; Suarez 2016). It is unclear if sperm grouping within an ejaculate is random or if certain cells (e.g., slower sperm) tend to group together, thereby forming slower aggregates. Moreover, because the viscosity female reproductive fluids in *Peromyscus* are unknown, it is not clear how they compare to our artificial suspension media or if they differ among species, thus we are limited in our interpretation of how sperm aggregation influences movement in the female reproductive fluids (Gasparini et al. 2020). In addition, a recent study reported that cell-cell interactions may be important for sperm migration through environments that mimic the highly-folded epithelium of the mammalian female reproductive tract (Bukatin et al. 2020). We can say with confidence, however, that sperm aggregates are not always faster than solitary cells, nor do they necessarily always perform better in viscous environments. Overall, we observed that sperm cells, including sperm aggregates, are slower as the viscosity of their environment is increased.

In only the promiscuous *P. maniculatus* did we find that sperm aggregates are relatively faster than solitary sperm in every environmental context regardless of viscosity, supporting the functional hypothesis that sperm aggregates evolved to be faster in the face of sperm competition (Moore et al. 2002; Immler et al. 2007). Its monogamous congener, *P. polionotus*, produces similarly large sperm aggregates, which are significantly bigger than the aggregates of other species, but *P. polionotus* sperm aggregates are slower in low viscosity media and no faster than solitary cells in more viscous environments. Together these findings suggest that the emergence of sperm aggregation likely evolved prior to the divergence of these two present-day species, which have been under divergent selective pressures due to differences in mating systems (Bedford and Hoekstra 2015). Our findings support that relaxed sexual selection and lack of intense sperm competition in *P. polionotus* has allowed for the degeneration of sperm aggregate motility (van der Horst and Maree 2014), despite the persistence of high levels of sperm aggregation. Given that *P. polionotus* aggregates were no faster than solitary sperm in any of the experimentally viscous environments, this suggests a loss of functional significance, although we cannot rule out a different function for sperm aggregation in this species that we failed to explore in this study. For example, agglutinated bovine sperm have increased motility and more functional mitochondria, suggesting that they maintain viability and motility for a longer period of time *in vitro*, thus may be important in mammalian fertilization (Umezu et al. 2020).

Our finding that the relative motility advantages of sperm aggregation differs depending on the environment for the other focal *Peromyscus* species corroborates studies in more disparate taxonomic groups, which have quantified collective sperm motility and found inconsistent results. For example, sperm trains exhibit greater progressive motility in the wood mouse (Moore et al. 2002), and greater velocity than single sperm in the Norway rat, but not the house mouse (Immler et al. 2007). In invertebrates, the swimming velocity of fishfly sperm increases with number of sperm in a bundle (Hayashi 1998), but in a marine snail, there is no differences in swimming speed between paired and single sperm (Ishijima et al. 1999). Although it was beyond the scope of this study, additional work is needed to elucidate the conditions within the female reproductive tract (e.g., fluid viscosity) so that they can be approximated *in vitro*, or observed *in vivo* (Ishikawa et al. 2016; Wang and Larina 2018; Qu et al. 2020), to determine if sperm aggregation does indeed allow for more efficient migration through viscous female secretions (Moore and Taggart 1995; Suarez 2016).

Sperm cells are predicted to be faster if they generate increased force with proportionally less drag; such effects may also be true for sperm aggregations in which cells conjoin head-to-tail, thereby increasing the length of the collective unit (Higginson and Pitnick 2011). In *Peromyscus*, sperm conjoin head-to-head to form aggregates (Fisher et al. 2014) and thus are wider, not longer, yet can still offer a motility advantage in some instances. This kinematic benefit, however, was not observed in all species, which led us question whether aggregate motility is dependent on the orientation of the cells within a group, or the inclusion of immotile or morphologically abnormal sperm (e.g., kinked or broken cells). When sperm heads and flagella are not oriented in the same direction, the cells within an aggregate are expected to exert opposing forces on one another, thereby reducing the overall motility of the group (Fisher et al. 2014; Pearce et al. 2018). Similarly, the inclusion of immotile or morphologically abnormal sperm, which typically display reduced motility (reviewed in Hook and Fisher 2020), is expected to reduce overall group kinematics. Indeed, we found that aggregates with misaligned, immotile and/or morphologically abnormal sperm were slower than aggregates without these abnormalities in all but one species. Perhaps not surprisingly, the species with the fewest abnormal aggregates, the promiscuous deer mouse (*P. maniculatus*), also showed the greatest kinematic improvement when sperm formed groups. These findings provide evidential support for the hypothesis that the orientation of sperm within the group is critical to its kinematics (Fisher et al. 2014) and that, in addition to uniformity of aggregate size, post-copulatory sexual selection may additionally act upon the architecture of sperm aggregates to improve their motility in a competitive context.

In contrast to our findings in *P. maniculatus*, sperm from their monogamous sister-species, *P. polionotus*, often forms aggregates that contain misaligned, immotile or morphologically abnormal cells, and overall *P. polionotus* aggregates are slower or no different in their speed than single cells. This finding is consistent with a previous report that *P. polionotus* sperm form optimal-sized aggregates less often than in *P. maniculatus* sperm (Fisher et al. 2014). Together, these observations are consistent with the theory that sperm aggregation evolved prior to the divergence of this species pair, and when monogamy evolved secondarily in *P. polionotus* (Greenbaum et al. 1978; Turner et al. 2010), relaxed sexual selection allowed for the persistence of less motile sperm traits (van der Horst and Maree 2014). In an examination of sperm conjugate presence or absence in 42 species of carabid beetles across nine subfamilies, results of the ancestral state reconstruction showed that sperm conjugation is ancestral and that the loss of conjugation evolved independently in several lineages (Sasakawa 2020). While sperm conjugation traits were not linked to mating systems per se, they do suggest that the evolutionary loss of these characters is prevalent. Based on our findings, we suggest that relaxed sexual selection may allow the persistence of less optimal sperm traits in the *Peromyscus* system.

In conclusion, our study highlights the diversity of sperm aggregation within a single taxonomic lineage and how post-copulatory sexual selection has likely shaped the formation and performance of these cellular groups. We show that in species expected to experience higher levels of sperm competition, sperm aggregates are less variable in size and males produce a fewer number of aggregates with inefficient alignment or composition. Moreover, we find that sperm aggregation can improve sperm motility, but this is not consistent across all species or contexts. Theoretical predictions (Fisher et al. 2014; Pearce et al. 2018) suggests that motility benefits may only be realized if cells maintain optimal alignment within the groups and, if achieved, collective motility may provide these sperm with a competitive advantage in the female reproductive tract (Higginson et al. 2012). Future work investigating *Peromyscus* sperm aggregates *in vivo* will help shed light on the co-evolution of collective sperm behaviors and the complex female reproductive tract through which sperm must navigate.

