## Supplementary Materials for "Post-copulatory sexual selection is associated with sperm aggregate quality in *Peromyscus* mice"

### Supplementary Methods

#### *Experimental animals*

While most of the focal males used in our experiment were 120-day old virgins, some individuals were older and/or were paired with a female prior to the start of the experiment. For example, we found that *P. eremicus* needed to be paired with a female and/or older than 120 days of age in order to actively produce sperm. In addition, we used several *P. californicus* males in our study that were simultaneously being used in another experiment for which those males had been previously paired with a female and were older. Hence, the mean (±SE) age of males used in our study was 193 (±12) days of age. After removing four clear outliers for sperm counts based on our hemocytometer estimates (see methods below) and based on non-normality of our model residuals when they were included (determined using a Shapiro test and visualized using ggplots of our model residuals), we found a significant effect of age on sperm counts (LM: $F_{1,117} = 5.92$, $p = 0.0165$). We found no such effect of pairing status (paired $n$ =32, unpaired $n$ =99, unknown $n$ =4) on sperm counts, however (LM: $F_{1,113} = 1.022$, $p = 0.314$). As a precaution, we considered both male age and pairing status as fixed factors in our statistical analyses for sperm aggregate size.

#### *Optimizing live sperm collections and observations*

A series of pilot experiments allowed us to pre-establish approaches that would reduce impacting sperm density and aggregate formation. These included the use of plastic rather than glass materials to reduce cells sticking to the slide surface (Fisher and Hoekstra 2010), imaging spacers with an adequate chamber depth (120μm) so as not to constrain sperm movement, pre-cut pipette tips to reduce breaking up already formed aggregates, as well as slow and gentle pressure and use of a reverse pipetting technique to avoid creating bubbles in the media and disperse sperm evenly throughout the imaging spacer. Our pilot experiments also allowed us to establish the following optimal CASA system parameter settings when recording videos of live *Peromyscus* sperm cells: head minimum brightness = 165, head size max ($\mu m^2$) = 475, head size min ($\mu m^2$) = 3, tail brightness min = 100, capillary correction = 1.0, chamber depth (μm) = 120, chamber type = drop.

Prior to our live cell video recording of sperm aggregates, we standardized the concentration of sperm cells across all samples by rapidly determining the concentration of sperm cells within each sample and performing a series of dilutions as needed. First, we pipetted 3ul of supernatant (i.e., collected from just below the meniscus) from the sperm cell sample into a 20μm deep chamber slide (SC- 20-01-04-B, Leja, The Netherlands) and observed the sample under an upright microscope with phase contrast (Axio Lab.A1, Zeiss, Germany) using a green filter (Hamilton Thorne, product number 720541) to enhance image contrast at 100X magnification. Using the CASA system, we recorded five 5-second videos in quick succession at multiple distances away from the site of entry for the loading chamber using the following CASA parameter settings: head brightness min = 150, head size max ($\mu m^2$) = 475, head size min ($\mu m^2$) = 1, tail brightness min = 50, capillary connection = 1.3, chamber depth (μm) = 20, chamber type = capillary, max photometer = 70, min photometer = 65, progressive STR (%) = 80, progressive VAP (μm/s) = 50, slow VAP (μm/s) = 20, slow VSL (μm/s) = 30, static VAP (μm/s) = 4, static VSL (μm/s) = 1, frame capture speed (Hz) = 60, frame count = 100. The CASA-generated analysis of these video recordings provided a summed count of cells as well as an estimate for total sperm concentration within each sample. We also manually estimated sperm density from these same samples using a Neubauer-improved hemocytometer (Marienfeld Superior, Germany), which is more time-consuming but is a more established and reliable technique (Rijsselaere et al. 2003; World Health Organization 2010). To do so, we collected an aliquot of the sample from just below the meniscus at the same time that we collected a sample for our CASA estimate, immediately fixed the cells by gently mixing the sample with formalin (SF98-4, 10% w/v, Fisher Scientific, USA) in a 9:1 ratio (e.g., 45μl of sperm to 5μl of formalin), and then loaded two replicate 10μl aliquots of the homogenized solution onto the hemocytometer. We allowed the cells to settle to the bottom for several minutes before collecting images using Zen Lite 2.3 (Zeiss, blue edition, 2011) and a camera attached to a light microscope (Axioplan, Zeiss, Germany) at 400x magnification. We manually counted sperm cells from these images for each replicate of five square grids on the hemocytometer and took an average of the two estimates to determine the sperm concentration for each sample. From these data, we were able to verify that these manual sperm estimates strongly positively correlates with those rapidly assessed using CASA (LM: $F_{1,117} = 32.25$, $p < 0.001$). Four outliers were removed and the data were log-transformed to meet model assumptions of normality, although whether or not these outliers were removed yielded the same result.

Once we verified our CASA estimates could provide a quick and accurate assessment of cell concentration, we then used these estimates to carry out dilution procedures to reach the optimal sperm concentration for each sample prior to our live cell observations, which we determined was 60-80 cells per video (i.e., 300-400 cells per male sample across all videos) at 100X magnification for optimal and efficient cell tracking. To achieve this standard concentration of cells, we diluted samples assessed to be above this range with pre-warmed media and then re-assessed their sperm concentration using the above protocol until the optimal concentration was reached. Because we recorded our dilution procedures, we were able to use our hemocytometer estimates to verify that our dilutions did indeed work and made sperm concentrations approximately equal across males (sperm cell density *before* dilutions: LM: $F_{5,111} = 13.87$, $p = < 0.001$; sperm cell density across species *after* dilutions: LM: $F_{5,112} = 2.607$, $p = 0.02861$; but note that $p > 0.05$ for all pairwise species comparisons except for *P. maniculatus*-*P. eremicus* $p = 0.0239$). We removed four and three clear outliers for these statistical models respectively and log-transformed data for the '*before* dilutions' model to meet assumptions of normality (determined quantitatively using a

Shapiro test as well as visually using diagnostic ggplots to assess model residuals), although doing so did not change the results. Post-hoc pairwise comparisons were conducted using the "LSmeans" R package (Lenth 2016). Because some samples fell below ($n$ = 40) and above ($n$ = 21) our ideal sperm concentration, we considered cell density ('total sperm cells') as a covariate within our statistical analyses of sperm aggregate size.

*Sperm observations in high-viscosity media*

To produce high-viscosity media to assess sperm aggregate kinematics in a complex environment, we warmed up an centrifugre tube containing 1.66 mL of modified sperm washing medium (MSWM) in an incubator (Incubator Genie, model SI-1400, Scientific Industries, Inc) to 75ºC, at which point we added 0.2g of methylcellulose to it. A series of steps were required to dissolve the methylcellulose, including agitating the sample for five minutes in the warmed incubator, adding 3.33 mL of cold MSWM into the tube and then agitating it for another five minutes in the warmed incubator, placing the sample in a 4ºC refrigerator for 20 minutes, and then agitating the sample at room temperature for another 30 minutes (Hyun et al. 2012). This protocol produced a 4% methylcellulose solution, which we then aliquoted into smaller centrifuge tubes containing 1 part of the 4% methylcellulose solution to 3 parts of MSWM (not enriched with methylcellulose) to produce a 1% methylcellulose media solution. We used the same protocol to create a 2% methylcellulose solution containing 2 parts of the 4% methylcellulose solution to 2 parts of MSWM as well as a 3% methylcellulose solution containing 3 parts of the 4% methylcellulose solution to 1 part of MSWM. We vortexed these solutions to homogenize them and then stored the tubes in a -20ºC freezer until use. Prior to use, each tube containing 60µl of the suspended methylcellulose solution was dethawed, vortexed again, and then warmed to 37ºC. Once we mixed in the sperm supernatant, our final solutions for the high-viscosity treatment groups were a 0.75%, 1.5%, and 2.25% methylcellulose media (made from 1%, 2%, or 3% methylcellulose tubes, respectively). The more methylcellulose suspended in media, the more viscous the solution.

*Statistical Analyses*

To investigate differences in sperm aggregate size across our focal *Peromyscus* species while including the single outlier *P. californicus* male, we used the same statistical methods outlined in the main manuscript, resulting in the use of a LM rather than a LMM; however, the response variable had to be log transformed to meet model assumptions of normality and heteroskedasticity given the single large outlier.

### Supplementary Results

When we include the outlier *P. californicus* male in our dataset, we found that sperm aggregate size still significantly differs among *Peromyscus* species (LM: $F_{6,127} = 46.31$, $P < 0.001$), indicating substantial trait variation within this lineage. We found greater variance in the number of aggregated cells between than within species (proportion: $s^2$ across species = 1.96; $s^2$ within species < 1.00, except for *P. californicus* [$s^2 = 2.90$] and *P. polionotus* [$s^2 = 2.15$]). Similar to our other analysis, the largest aggregates were observed in sister species *P. maniculatus* and *P. polionotus*, which did not statistically differ from one another in their size ($P = 0.81$). Pairwise comparisons adjusted for multiple comparisons using LSmeans revealed that *P. leucopus*, *P. eremicus*, *P. gossypinus*, and *P. californicus* all produce similarly sized smaller aggregates ($P > 0.05$ for pairwise comparisons), with the exception of P. *californicus* and *P. gossypinus*, with the latter producing statistically smaller aggregates than the former ($P = 0.047$).

When we included the outlier *P. californicus* male in our dataset, the CV for sperm aggregate size within *P. californicus* was 54.3%. Using a PGLS regression, we found the same negative association between mean relative testis weight and sperm aggregate size CV (PGLS: $F_{2,3} = 8.768$, $t = -3.1520$, $p = 0.05$, $r = 0.76$, $\lambda = 0^{1,0.01}$) but no such association with mean sperm aggregate size (PGLS: $F_{2,3} = 2.836$, $t = -2.0073$, $p = 0.1384$, $r = 0.42$, $\lambda = 0^{1,0.51}$).

**Supplementary Figures and Tables:**

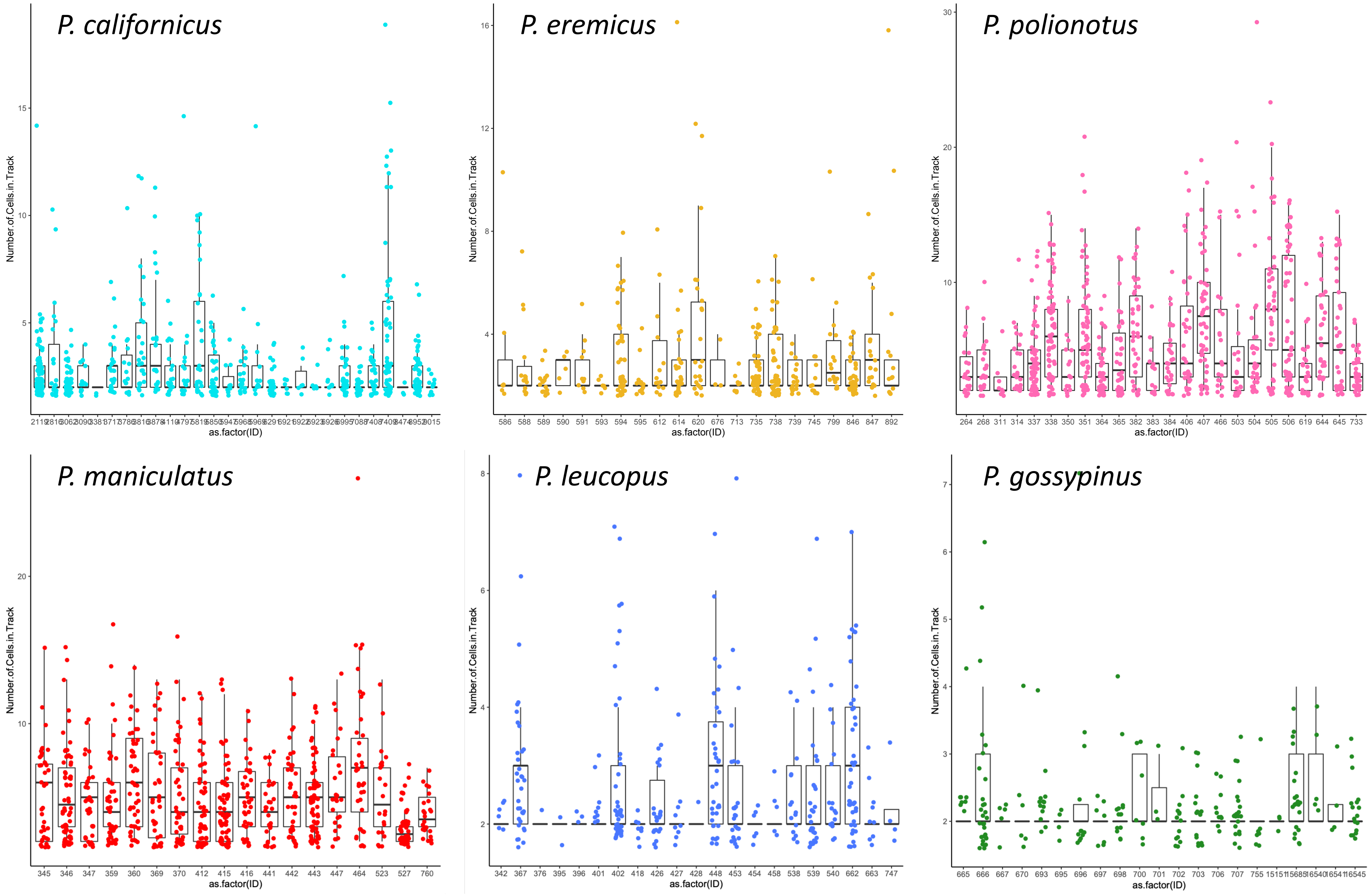


**Figure S1.**

The observed intra- and inter-male variation in the number of sperm cells per aggregate naturally produced by individual males for six species of *Peromyscus* rodents under the same controlled *in vitro* conditions. Box-plots represent median and interquartile ranges with mean values per male overlaid as dots.

**TABLE S1.**

Results from an intra-male analysis for six species of *Peromyscus* mice comparing the curvilinear velocity of solitary and aggregated sperm cells suspended in media enriched with varying amounts of methylcellulose to test whether sperm aggregates confer a kinematic advantage in more viscous environments (shaded in gray)

| *Peromyscus* Species | 1.5% Solution | | | | | 2.25% Solution | | | | |
|---|---|---|---|---|---|---|---|---|---|---|
| | Mean (±SE) curvilinear velocity (µm/sec) | | Student's paired t-test | | | Mean (±SE) curvilinear velocity (µm/sec) | | Student's paired t-test | | |
| | Solitary cells | Aggregated cells | *t* | df | *p* | Solitary cells | Aggregated cells | *t* | df | *p* |
| *P. californicus* | 59.1 ± 4.9 | 70.3 ± 6.2 | -3.137 | 5 | **0.026** | 45.5 ± 4.3 | 58.5 ± 5.1 | -3.164 | 11 | **< 0.01** |
| *P. eremicus* | 51.1 ± 4.7 | 56.3 ± 5.4 | -2.422 | 7 | **0.046** | 54.5 ± 4.9 | 57.9 ± 8.8 | -1.179 | 7 | 0.2769 |
| *P. polionotus* | 71.8 ± 4.9 | 72.9 ± 4.4 | -0.707 | 12 | 0.493 | 59.6 ± 5.0 | 66.5 ± 5.0 | -2.156 | 12 | 0.052 |
| *P. maniculatus* | 75.1 ± 3.9 | 88.2 ± 5.0 | -5.2835 | 11 | **< 0.001** | 67.2 ± 5.4 | 82.7 ± 5.2 | -5.3274 | 11 | **< 0.001** |
| *P. leucopus* | 71.0 ± 3.8 | 72.6 ± 5.4 | -0.506 | 8 | 0.627 | 59.0 ± 5.5 | 76.2 ± 5.0 | -2.450 | 7 | **0.044** |
| *P. gossypinus* | 68.8 ± 3.7 | 74.9 ± 3.8 | -2.783 | 5 | **0.039** | 69.0 ± 4.2 | 64.2 ± 4.6 | -0.696 | 5 | 0.517 |

Significant values at the $p < 0.05$ level are bolded.
